# Utility of cell-free DNA concentrations and illness severity scores to predict survival in critically ill neonatal foals

**DOI:** 10.1101/2020.11.09.373944

**Authors:** Sarah F. Colmer, Daniela Luethy, Michelle Abraham, Darko Stefanovski, Samuel D. Hurcombe

## Abstract

Plasma cell-free DNA (cfDNA) levels have been associated with disease and survival status in septic humans and dogs. To date, studies investigating cfDNA levels in association with critical illness in foals are lacking. We hypothesized that cfDNA would be detectable in the plasma of foals, that septic and sick-nonseptic foals would have significantly higher cfDNA levels compared to healthy foals, and that increased cfDNA levels would be associated with non-survival. Animals used include 80 foals of 10 days of age or less admitted to a tertiary referral center between January and July, 2020 were stratified into three categories: healthy (n=34), sick non-septic (n=11) and septic (n=35) based on specific criteria. This was a prospective clinical study. Blood was collected from critically ill foals at admission or born in hospital for cfDNA quantification and blood culture. Previously published sepsis score (SS) and neonatal SIRS score (NSIRS) were also calculated. SS, NSIRS, blood culture status and cfDNA concentrations were evaluated to predict survival. Continuous variables between groups were compared using Kruskal-Wallis ANOVA with Dunn’s post hoc test. Comparisons between two groups were assessed using the Mann-Whitney *U*-test or Spearman rank for correlations. The performance of cfDNA, sepsis score and NSIRS score to predict survival was assessed by receiver operator characteristic (ROC) curve analysis including area under the curve, sensitivity and specificity using cutoffs. Plasma cfDNA was detectable in all foals. No significant differences in cfDNA concentration were detected between healthy foals and septic foals (*P*=0.65) or healthy foals and sick non-septic foals (*P*=0.88). There was no significant association between cfDNA and culture status, SS, NSIRS or foal survival. SS (AUC 0.85) and NSIRS (AUC 0.83) were superior to cfDNA (AUC 0.64) in predicting survival. cfDNA was detectable in foal plasma however offers negligible utility to diagnose sepsis or predict survival in critical illness in neonates.

## INTRODUCTION

Sepsis is the most common cause of neonatal foal mortality and is broadly defined by an overwhelming systemic inflammatory response syndrome triggered by infectious organisms [1]. A myriad of factors are involved in the pathophysiology of neonatal sepsis, including maternal disease, parturition complications and management practices [2]. The clinical signs observed in sepsis overlap with many other non-infectious, life-threatening conditions, and due to this ambiguity, diagnosis proves difficult until the disease become severe [3]. Recent studies show septic foals still have a high mortality rate of 30-50% despite improved survival with medical advances [4,5].

The gold standard for the detection of bacteremia is blood culture which exhibits low sensitivity in neonates at approximately 61.9% [6]. False negative blood cultures in septic foals as well as positive blood cultures documented in healthy foals, further complicate the interpretation of results [7,8]. Blood cultures regularly require up to one week before they can be considered diagnostic. Scoring systems were developed in the 1980s using blood culture and clinical findings to expedite and predict a diagnosis of sepsis in foals. These scoring systems incorporated both subjective and objective clinical and clinicopathologic findings and have yielded varying diagnostic accuracy suggesting variation within at-risk foal populations at different times and locations [9].

Recent studies in human and veterinary medicine have shown the prognostic value of cell-free DNA (cfDNA) in the blood of septic patients [10–12]. cfDNA is proposed to be released from various cell types during apoptosis and necrosis, and thought to be involved in host defense systems with the intent to trap and kill bacteria [13]. Previously, quantification of cfDNA was a labor- and cost-intensive process involving DNA isolation, extraction, gel electrophoresis and real-time PCR [14]. More recently, rapid and direct fluorescence assays have been developed that use nucleotide-binding fluorophores to identify cfDNA directly within biological fluids [14–16].

In humans, cfDNA concentrations have proven to be sensitive and specific for poor outcomes [17,18]. One study analyzing 255 patients with severe disease revealed that admission cfDNA concentrations were higher in ICU non-survivors than in survivors [18]. A 2017 study assessing cfDNA concentrations in dogs using fluorometric assays found significantly higher concentrations in those presenting to an emergency service with sepsis or non-septic SIRS compared to healthy dogs [19]. Though the body of cfDNA research is growing in both human and veterinary medicine, there are no studies documenting its utility in equine neonatal sepsis to the authors’ knowledge.

We believe that cfDNA may have diagnostic and prognostic utility for diagnosis of sepsis in critically ill foals. We hypothesized that cfDNA would be detectable in the plasma of newborn foals and that the magnitude of cfDNA concentrations would be proportional to the severity of illness. Our specific aims were to measure and compare cfDNA from healthy, sick non-septic and septic foals at admission. An additional aim was to determine if there is an association between cfDNA concentrations and detectable bacteremia as well as previously published critical illness scoring systems, namely sepsis score and neonatal systemic inflammatory syndrome score [20]. Finally, we sought to determine if cfDNA concentrations could predict survival in critically ill foals.

## MATERIALS AND METHODS

### Study Population

The study population consisted of foals ≤ 10 days who were admitted to or born at the University of Pennsylvania School of Veterinary Medicine New Bolton Center between January and July 2020. Inclusion criteria required the results of a blood culture performed within 24 hours of admission, documentation of either discharge from the hospital or non-survival, and recorded physical examination and clinicopathological parameters for the implementation of the updated sepsis scoring system and neonatal SIRS (NSIRS) scoring system published by Wong et al. in 2018 [20]. Foals were excluded if they were > 10 days of age and did not have a blood culture performed during hospitalization. All experimental procedures were approved by the University of Pennsylvania Institutional Animal Care and Use Committee (protocol # 806836) and undertaken with informed client consent.

### Classification

Foals were classified into 1 of 3 groups: septic, sick non-septic and healthy [21,22]. Foals were defined as septic if they fulfilled any of the following criteria: (1) positive blood culture, and/or a sepsis score of ≥ 12 [20]. Sick non-septic foals were defined as those with a negative blood culture and a sepsis score between 6 and 11. Healthy foals were defined as those with a negative blood culture and a sepsis score ≤ 5.

Survival was defined as any foal admitted to New Bolton Center who was discharged alive. Non-survival was defined as any foal born or admitted to New Bolton Center who died or was euthanized due to grave medical prognosis. Foals euthanized due to financial constraints were excluded.

### Data Collection

Historical data including maternal health during pregnancy, foaling date and details of foaling (Cesarean section, dystocia, induction protocol) were obtained. Age at presentation, breed and sex were recorded. Physical examination findings including rectal temperature, heart rate, respiratory rate, the presence of swollen joints, diarrhea, petechiation, scleral injection, hypopyon, anterior uveitis, respiratory distress and the presence of seizures were recorded. The sepsis score (SS) and neonatal systemic inflammatory response syndrome score (NSIRS) were calculated for each foal [20]. Outcome was recorded as survival to hospital discharge or non-survival.

Clinicopathologic data collected included a compete blood count (CBC) (Element HT5, Heska, Loveland, CO), serum biochemistry (VITROS 350 Chemistry System, Ortho Clinical Diagnostics, Buckinghamshire, England), blood glucose concentration (Accu-Chek Performa glucometer, Roche, Indianapolis, IN), blood L-lactate concentration (Nova 8+ Electrolyte analyzer, Nova Biomedical, Waltham, MA), plasma fibrinogen concentration (ACL Elite, Instrumentation Laboratory, Bedford, MA), serum immunoglobulin G (IgG) concentration (DVM Rapid Test, MAI Animal Health, Elmwood, WI), blood culture (BD SEPTI-CHEK media bottles, BD Biosciences, Franklin Lakes, NJ) (VersaTREK Automated Microbial Detection System, ThermoFischer Scientific, Pittsburgh, PA) and cfDNA concentration (Qubit 4 fluorometer, Fisher Scientific, Pittsburgh, PA and Qubit™ dsDNA HS Assay Kit, Fisher Scientific, Pittsburg, PA).

### Sampling and processing

Approximately 25 mL of whole blood was obtained by jugular venipuncture using aseptic technique at the time of catheter placement or within 12 hours of admission or birth. Blood cultures were processed via inoculation (8mL of blood) and incubation of SEPTI-CHEK media bottles at 35°C for up to 7 days and detection using the VersaTREK Automated microbial system. When turbidity was apparent or 7 days had elapsed, the broth was Gram-stained and subcultured using trypticase soy agar with 5% sheep’s blood and MacConkey agar for aerobic isolation and using Brucella blood agar for anaerobic isolation.

Citrated plasma was collected for cfDNA quantification. Briefly, 2.7 mL of whole blood was collected into a citrate collection tube (BD Biosciences, Franklin Lakes, NJ) and centrifuged (Centritic Centrifuge, Fisher Scientific, Pittsburgh, PA) for 10 minutes at 1370 g within 30 minutes of collection. Plasma was then pipetted into polypropylene conical tubes. A small amount of plasma was left in the citrate tube so as not to disturb the buffy coat. Tubes were stored at −40°C, which has previously been shown to preserve sample integrity for cfDNA analysis [23]. At the time of batch processing, samples were thawed by allowing to come to room temperature (20-22°C) and cfDNA quantification measures were performed in triplicate using a benchtop fluorometer (Qubit 4 fluorometer, Fisher Scientific, Pittsburgh, PA) and associated reagents (Qubit 1x dsDNA HS Assay Kit, Fisher Scientific, Pittsburgh, PA) in thin-walled polypropylene tubes (Qubit Assay Tubes, Invitrogen, Carlsbad, CA) according to manufacturer’s instructions.

### Statistical analysis

Prior to initiating the study, sample size calculation was performed based on prior prevalence data in proportions of critically ill foals who were bacteremic [24]. With a power of 80% and significance set at *P* < 0.05, at least sixty (60) animals were deemed necessary for this prospective study. Data were assessed for normality using the Shapiro-Wilk statistic. Non-parametric data are represented as median and interquartile range or standard error of the mean unless otherwise stipulated. Continuous variables between groups were compared using Kruskal-Wallis ANOVA with Dunn’s post hoc test. Comparisons between two groups (blood culture positive and culture negative; survivor and non-survivor) were assessed using the Mann-Whitney *U*-test. Correlations between continuous variables were evaluated using Spearman rank test. The performance of cfDNA, sepsis score and NSIRS score to predict survival was assessed by receiver operator characteristic (ROC) curve analysis including area under the curve, sensitivity and specificity using cutoffs. Significance was set to *P* < 0.05. Analyses were performed using Prism Version 8.4.2 (GraphPad Software, San Diego, CA) and STATA (STATA Corp LLC, College Station, TX).

## RESULTS

### Study Population

A total of 80 foals were enrolled in the study between January and July, 2020. There were 34 healthy foals (43%), 35 septic foals (44%), and 11 sick non-septic foals (13%). Of the 80 foals, 45 were colts (56%) and 35 were fillies (44%). Breeds represented within the study population included Thoroughbred (n=48), Standardbred (n=18), Warmblood (n=7) and 1 each of the following: Appaloosa, Arabian, Clydesdale, Friesian, Miniature Horse, Morgan and Pony. There was no significant difference in sex or breed between the 3 groups. A total of 69/80 (86%) foals survived to discharge, and 11/80 (14%) of foals were euthanized or died. Of those that died, 5/11 (45%) had a positive blood culture. Of the 5 with positive blood cultures, 5/5 (100%) grew Gram-positive organisms, while 2/5 (40%) also grew a Gram-negative organisms. Isolates were of the following genus: *Bacillus*, *Acinetobacter*, *Micrococcus*, *Staphylococcus*, *Dietzia*, *Klebsiella* and *Enterococcus*.

Septic foals as defined using SS ≥ 12 and/or positive blood culture had a median sepsis score of 10 (range 0 to 29) at the time of presentation. The median NSIRS score for septic foals was 2 (range 0 to 5). In the septic group, the mortality rate was 26% (9/35), of which 2/9 (22%) died in hospital and 7/9 (78%) were euthanized. The most commonly diagnosed disease process/comorbidity at the time of admission in the septic foal group was neonatal encephalopathy (n=12), followed by diarrhea (n=5), failure of transfer of passive immunity (n=4), dystocia (n=3) and dysmaturity (n=3). Additional diagnoses included colic (n=1), dysphagia (n=1), atresia coli (n=1) and Lethal White Foal Syndrome (n=1).

Healthy foals, as defined by a sepsis score ≤ 5 and negative blood culture had a median sepsis score of 2.5 (range 0 to 5) and median NSIRS score of 1 (0 to 2) at the time of presentation. The mortality rate was 0% (0/34). The most common reasons for presentation at the time of admission was birth as part of a standard foaling program package for low-risk mares (n=11), mild neonatal encephalopathy (n=8) and meconium impaction (n=4). Three foals were presented as companions to their dams and two foals were presented for mild idiopathic dysphagia. Two foals were presented as part of a standard foaling program package for high-risk mares. Sick non-septic foals had a median sepsis score of 9 (range 6 to 11) and median NSIRS score of 1 (range 0 to 2) with a mortality rate of 22% (2/9). Both animals were euthanized due to grave prognosis. The most common reasons for presentation at the time of admission were dystocia (n=3) and diarrhea/enterocolitis (n=3).

Of foals that had positive blood cultures (n=24), 39 isolates were obtained. Of those isolates, 23/39 (59%) were Gram-positive and 16/39 (41%) were Gram-negative. The most common isolates were of the genus *Staphylococcus* (n=7), *Bacillus* (n=3), *Acinetobacter* (n=3), *Escherichia* (n-3) *Streptococcus* (n=2), *Microbacterium* (n=2), *Pseudomonas* (n=2), *Stenotrophomonas* (n=2), *Arthrobacter* (n=2), *Pantoae* (n=2) *and Enterococcus* (n=2).

Additional isolates (n=1) included *Actinobacillus*, *Rhizobium*, *Curtobacterium*, *Corynebacterium*, *Klebsiella*, *Weissella*, and *Dietzia*. The mean NSIRS score of the bactermic population was 1.64 (standard deviation = 1.15), and the mean NSIRS score of the non-bacteremic population was 1.26 (standard deviation = 1.21). The mean sepsis score of the bacteremic population was 8.29 (standard deviation = 7.01), whereas the mean sespsis score of the non-bacteremic septic population was 6.28 (standard deviation = 5.09).

### cfDNA concentrations and foal classification

There were no differences in cfDNA concentrations between groups of foals (**Fig 1**; P>0.05). Median cfDNA concentrations were 279.72 ng/mL (range 121.33 ng/mL to 424.67 ng/mL) in healthy foals, 301.33 ng/mL (range 193-393.67 ng/mL) in sick non-septic foals and 299.15 ng/mL (166.67 to 879.33 ng/mL) in septic foals. We further evaluated cfDNA concentrations based on SS alone to account for false positive blood cultures in healthy or non-septic foals and false negative blood cultures in septic foals. **Table 1** shows cfDNA concentrations between groups of foals based on SS cut-off criteria alone (septic ≥ 12, SNS 6-11, healthy ≤ 5) excluding blood culture results. cfDNA concentrations were not different between foals when classifying based on SS alone (healthy versus septic P=0.37; sick non-septic versus septic P=0.43).

**Figure 1:**
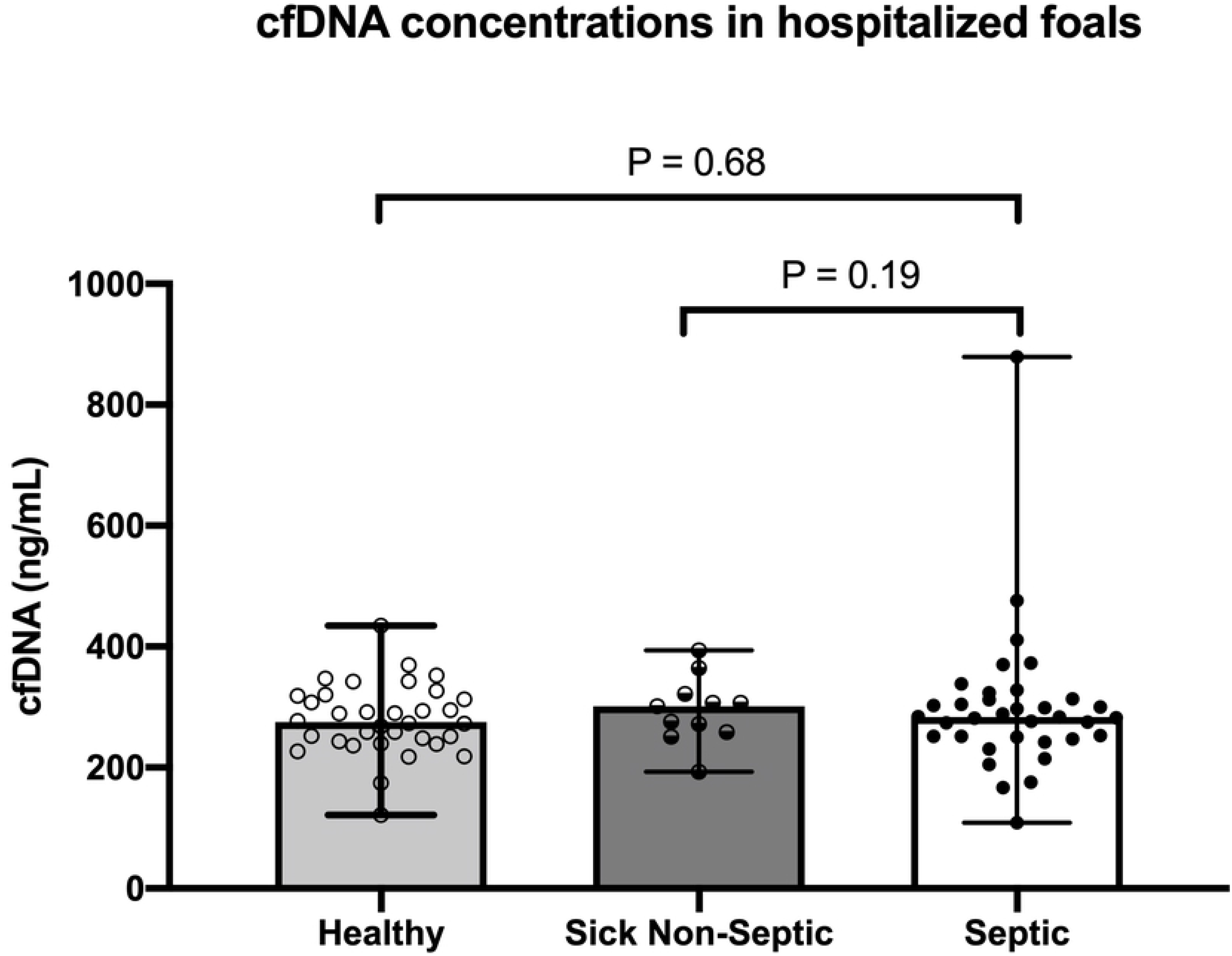
Plasma cell-free DNA (cfDNA) concentrations in healthy foals (n=34), sick non-septic foals (n=11) and septic foals (n=35) based sepsis score and/or blood culture status. Symbols represent values for individual foals with median and interquartile range error bars (P>0.05 between groups).

**Table 1:** Median (SE) cell-free DNA concentrations in 80 hospitalized neonatal foals based on sepsis score cut-off values alone, including 44 healthy foals, 21 sick non-septic foals and 15 septic foals.

| Foal Category | Median (SE) cfDNA (ng/mL) | 95% Confidence Interval |
| --- | --- | --- |
| Healthy (SS $\leq 5$)<br>(n= 44) | 283.28 (7.52) | 268.52 to 298.03 |
| **Sick Non-septic (SS 6-11)**<br>**(n=21)** | 281.17 (13.15) | 255.39 to 306.94 |
| **Septic (SS ≥ 12)**<br>**(n=15)** | 253.05 (33.10) | 188.19 to 317.92 |
| Values expressed as median (standard error) and 95% confidence interval. SS = sepsis score. |  |  |

The relationship between sepsis score and cfDNA concentration is illustrated in **Fig 2A** and was not significantly correlated (P=0.48). Similarly, there was no significant correlation between NSIRS score and cfDNA concentration (**Fig 2B**; P=0.54). There were no significant difference in median cfDNA concentrations between bacteremic foals (298.67 ng/mL, range 121.3 to 434.7 ng/mL) and non-bacteremic foals (275.33, range 109 to 879.3 ng/mL; P=0.07; **Fig 3**).

**Figure 2:**
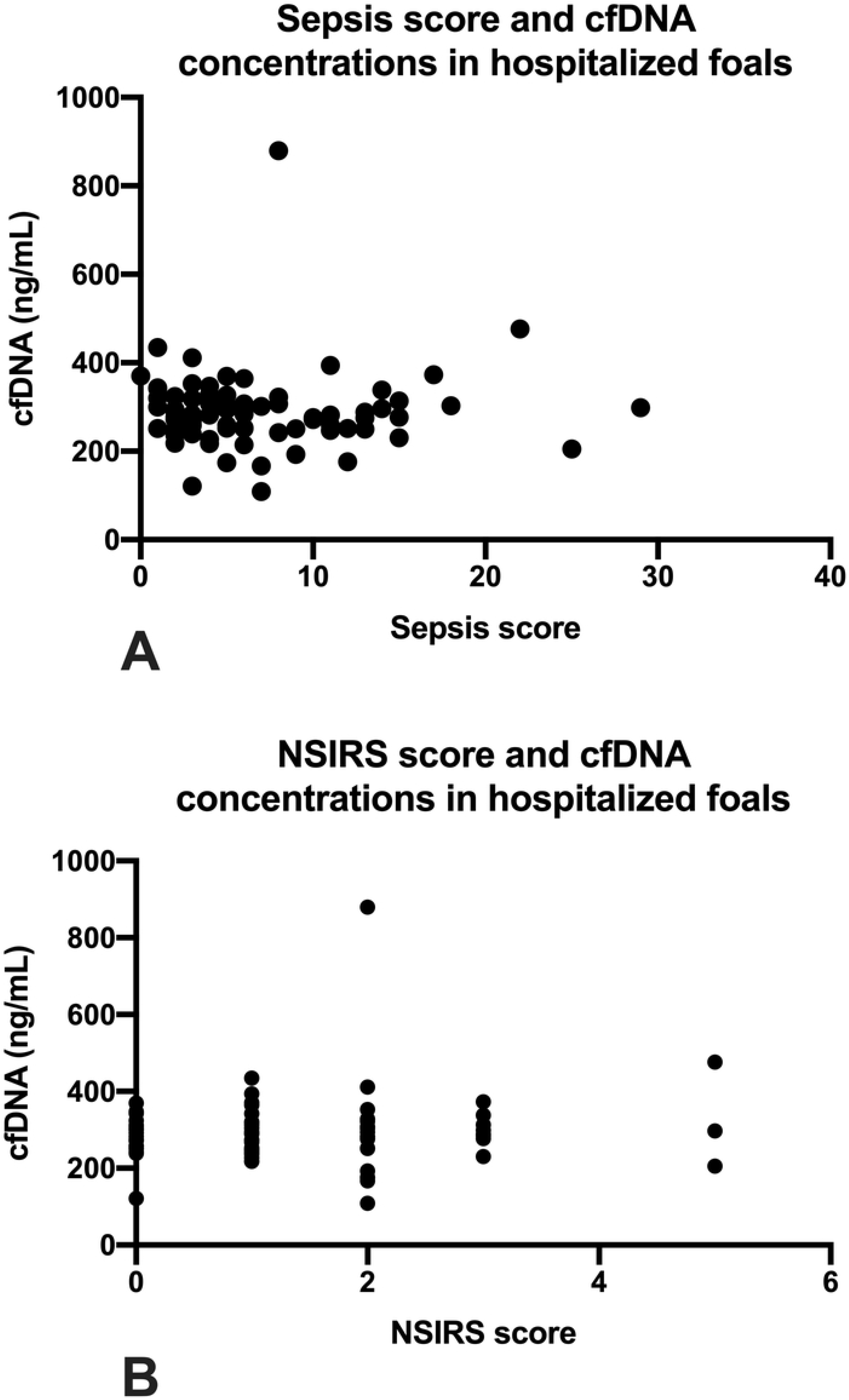
Plasma cell-free DNA (cfDNA) concentrations and illness severity scores in 80 hospitalized foals. 2A: cfDNA concentration and sepsis score (r= 0.07; P=0.48). 2B: cfDNA concentration and NSIRS score (r = 0.07; P=0.54). Symbols represent values for individual foals.

**Figure 3:**
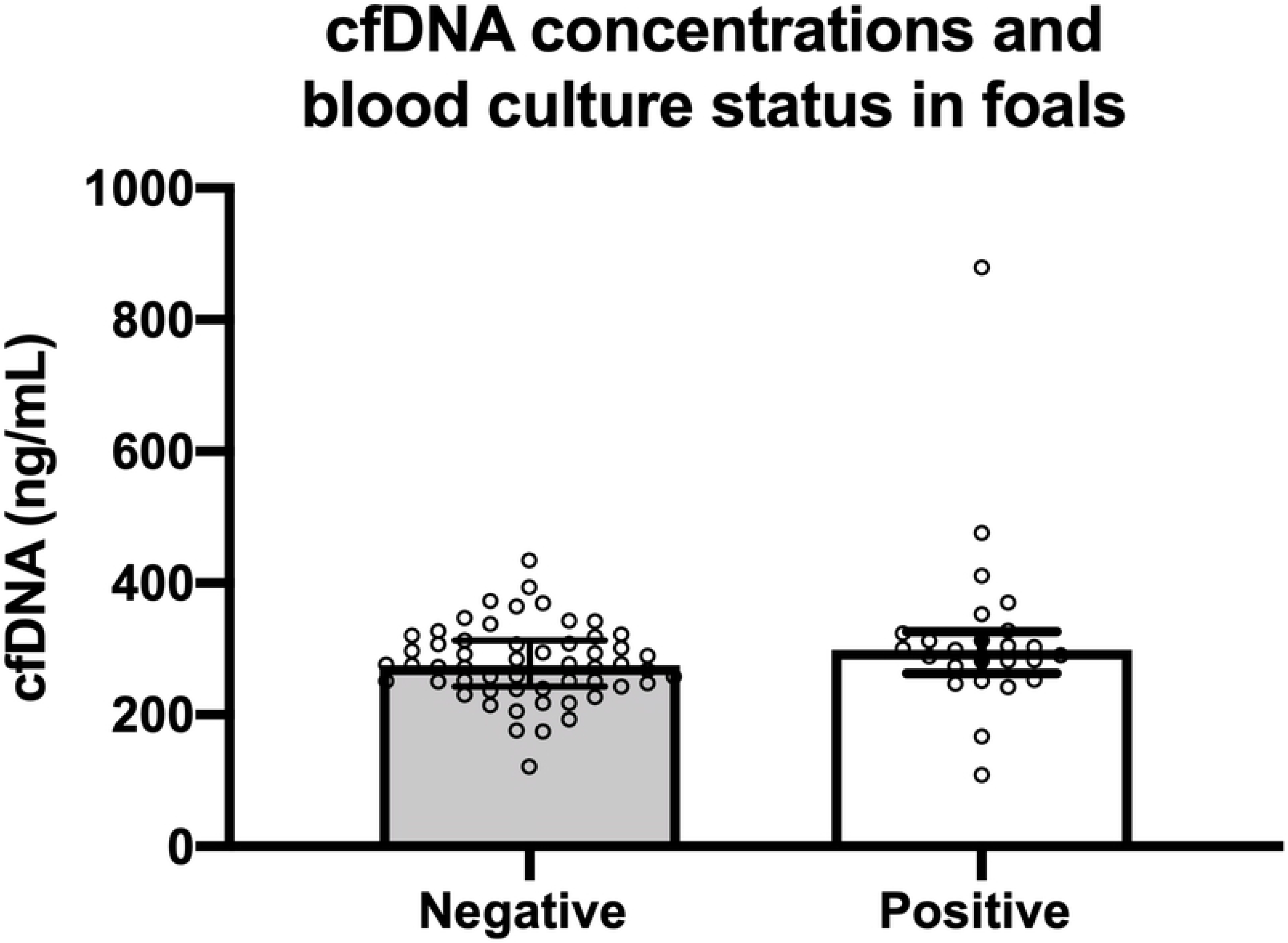
Plasma cell-free DNA (cfDNA) concentrations in 80 hospitalized neonatal foals with positive blood culture (n=24) and negative blood culture (n=56). Symbols represent values for individual foals with median and interquartile range error bars (P=0.07).

Further analysis evaluating the effect of age on cfDNA concentrations showed that for each day of age, there was a 1.72% increase in cfDNA concentration although not statistically significant (P=0.13).

### cfDNA concentrations and critical illness scoring systems in predicting outcome

cfDNA concentrations between survivors and non-survivors are shown in **Fig 4**. There was no significant difference in cfDNA concentrations between foals that survived (median 286.88 ng/mL (range 121.33 to 879.33 ng/mL) and non-survivors (314.5 ng/mL, range 176.00 to 476.33; P=0.16). Univariate logistic regression showed no association between survival and cfDNA concentration (P=0.41). Receiver operating characteristic (ROC) curve analysis revealed poor utility for cfDNA to predict survival (AUC 0.64, P=0.15; **Fig 5A**). ROC for sepsis score (**Fig 5B**) and neonatal SIRS score (**Fig 5C**) showed superior prediction to predict survival than cfDNA with an AUC for sepsis score of 0.85 (P<0.001) and for NSIRS score of 0.83 (P<0.001). **Table 2** shows cut-off values for cfDNA, SS and NSIRS to yield the optimal diagnostic accuracy for survival.

**Figure 4:**
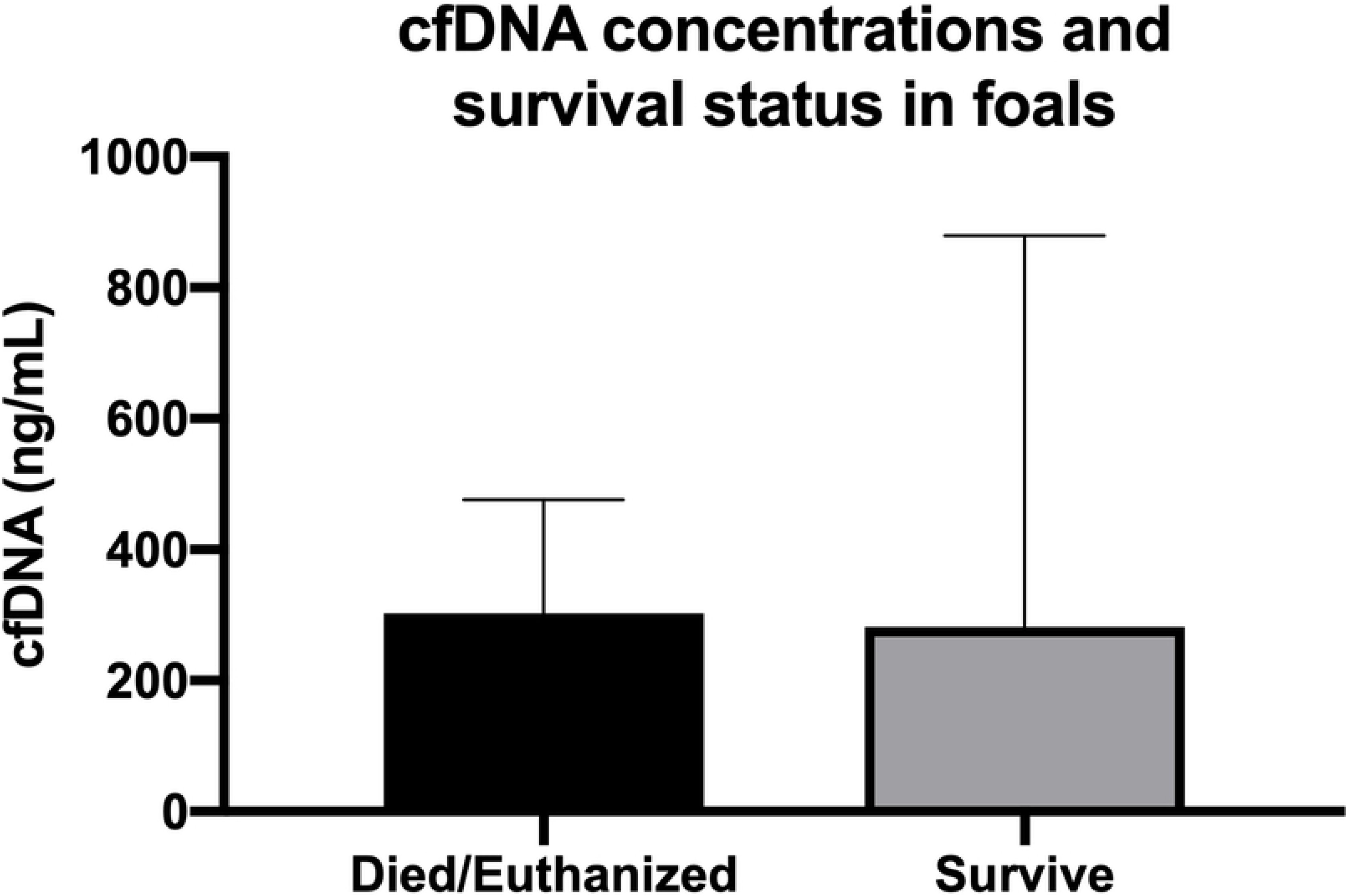
Plasma cell-free DNA (cfDNA) concentrations in 80 hospitalized neonatal foals that survived (n=69) and died (n=11). Symbols represent values for individual foals with median and interquartile range error bars (P=0.16).

**Figure 5:**
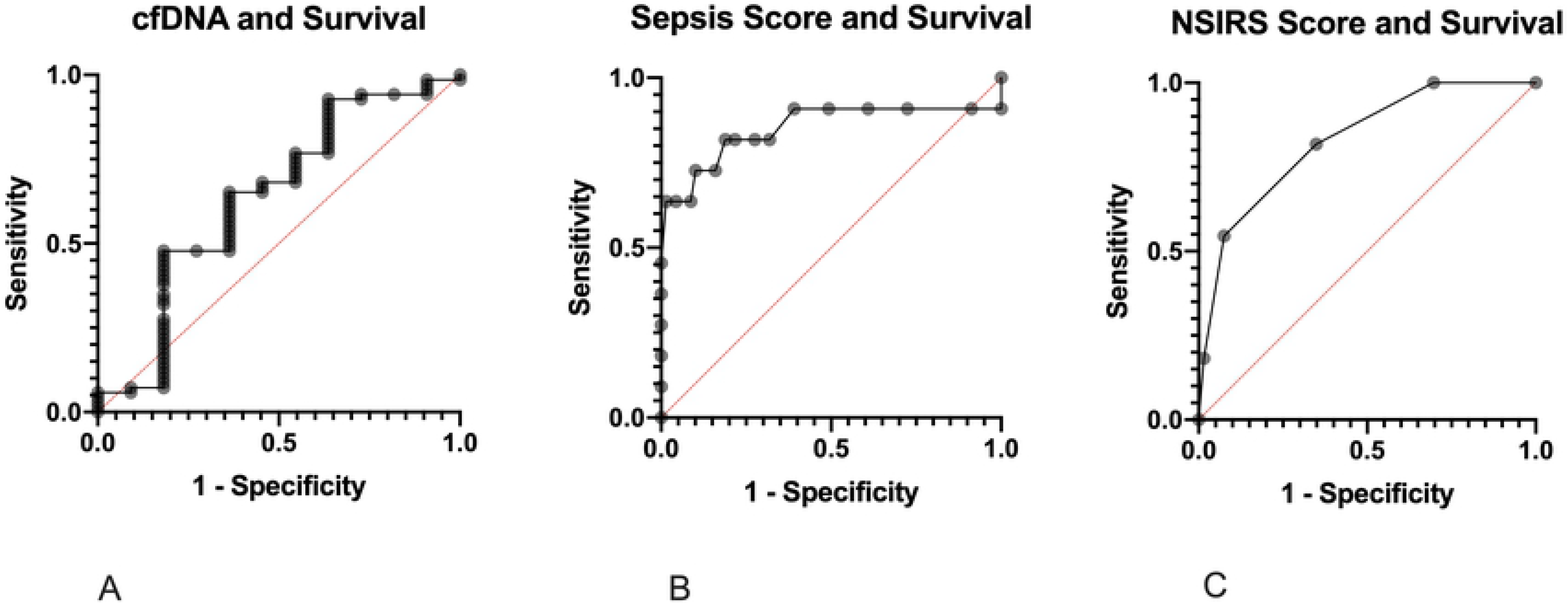
Receiver operator characteristic (ROC) curves for cfDNA (A), sepsis score (B) and NSIRS score (C) for predicting survival in 80 hospitalized neonatal foals.

**Table 2:**
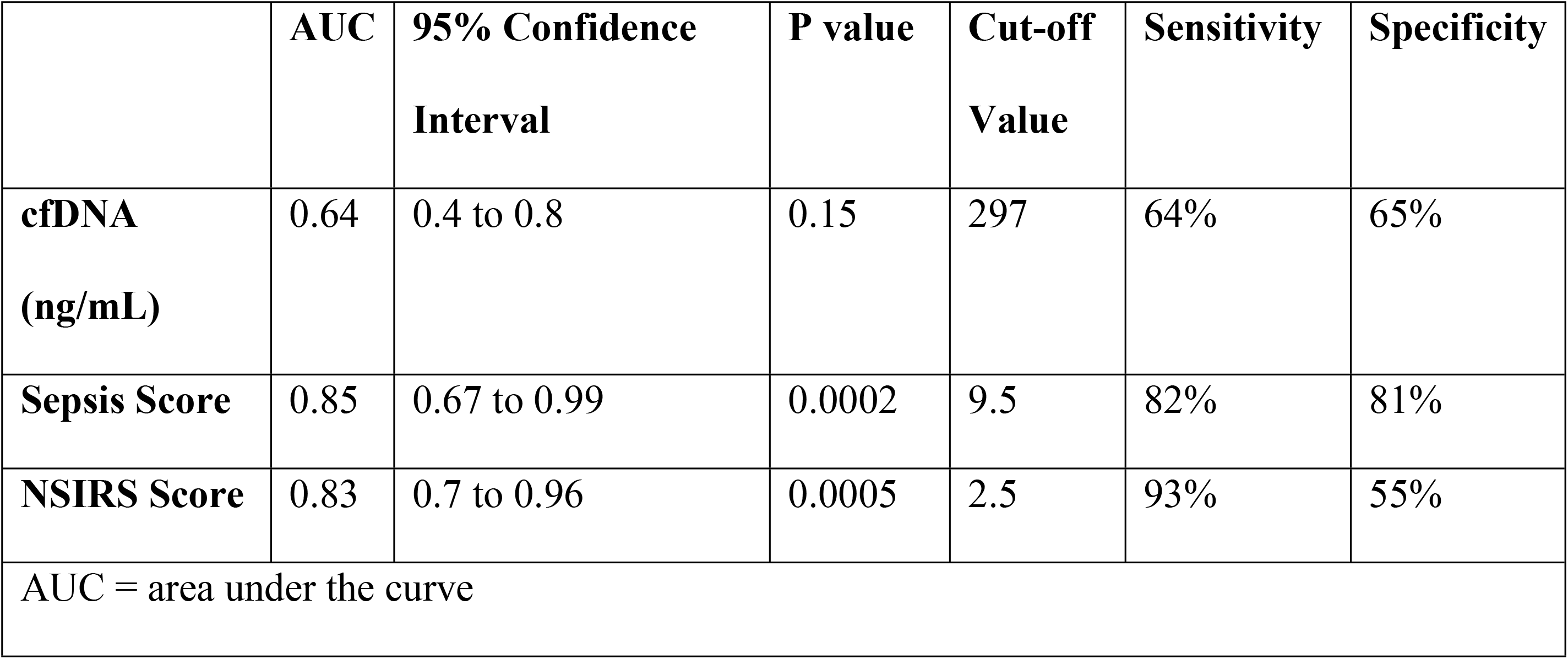
Diagnostic cut-off values for cfDNA, sepsis score (SS) and neonatal systemic inflammatory response syndrome (NSIRS) score to predict survival in 80 hospitalized neonatal foals.

|  | **AUC** | **95% Confidence**<br>**Interval** | **P value** | **Cut-off**<br>**Value** | **Sensitivity** | **Specificity** |
| --- | --- | --- | --- | --- | --- | --- |
| **cfDNA**<br>**(ng/mL)** | 0.64 | 0.4 to 0.8 | 0.15 | 297 | 64% | 65% |
| **Sepsis Score** | 0.85 | 0.67 to 0.99 | 0.0002 | 9.5 | 82% | 81% |
| **NSIRS Score** | 0.83 | 0.7 to 0.96 | 0.0005 | 2.5 | 93% | 55% |
| AUC = area under the curve |  |  |  |  |  |  |

## DISCUSSION

In the present study, cfDNA was detected in 100% of plasma samples obtained from this population of 80 foals admitted to a tertiary care facility. This finding supported our hypothesis that cfDNA would be detectable in plasma samples obtained from the equine neonate. Though circulating plasma cell-free DNA has been detected in multiple veterinary species including canines [10,19, 25,26] and felines [27], this is the first study confirming its presence in neonatal foals. Results of this study demonstrate a lack of significance in circulating cfDNA concentrations between foal cohorts with varying illness severity. Further, cfDNA concentrations were not correlated with SS or NSIRS and we did not demonstrate an association between cfDNA and blood culture status or survival.

Recent studies in humans [11,17,18,28] and dogs [10,19] have illustrated the value of cfDNA as a potential biomarker for the detection of sepsis through its ability to discriminate septic and healthy patients. Of note, all of these studies have been performed on adult populations, differing from the age group of the findings reported here. In 2008, Tuaeva et al. investigated levels of cfDNA in the blood plasma of premature human neonates [29]. The ranges of cfDNA concentrations obtained were between 9.9 and 136.5 ng/mL [29] across all groups, significantly lower than ranges in other studies performed in adults. This information combined with our observed trend of increasing cfDNA concentrations with age have led us to postulate that a reason for relatively low circulating cell-free DNA concentrations in foals could be a function of age and an immunonaive state [30].

Though there exists a lack of consensus on the origins of circulating cell-free DNA, it is believed that circulating levels are related cellular breakdown and active DNA-release mechanisms in host cells [31]. The concept of neutrophil extracellular traps, or NETs, involves the release of web-like scaffolds of cell-free DNA composed of extracellular chromatin which can enhance antimicrobial activity in the face of an infectious stimulus [32]. A model utilizing canine neutrophils has revealed that high-dose lipopolysaccharide stimulates the cells to undergo the production of NETs *in vitro* [33]. Investigating such mechanisms in the neonate, it is possible that our samples were obtained from an immune-competent but immune-naïve population based on age (10 days or less) where the degree of neutrophil activation, function or apoptosis is decreased compared to adult horses thereby limiting the amount of cell-free DNA released into circulation [34].

The relatively small number of foals that died or were euthanized in this study compared to previous reports may also have affected our ability demonstrate the utility of cfDNA to predict outcome [3,35]. In one published retrospective study analyzing equine neonatal admissions at a university teaching hospital between 1982 and 2008, 27.2% of foals did not survive to discharge [3]. A more recent retrospective of neonates admitted to a university or private referral hospital between 2008-2009 had a mortality rate of 21%. In this study, only 11/80 (14%) did not survive to discharge, limiting our ability to extrapolate the relationship between cell-free DNA levels and survival. It is possible that the difference between studies may lie in differences in patient populations and diseases, and the large number of healthy foals enrolled in the study may have contributed to these data. Further study with a larger population of critically ill foals may further define if cfDNA is associated with neonatal survival.

The definition of sepsis is a significant factor to consider in this study as well as past and future studies. This is particularly important in how foals are stratified and by what discriminating factors. Utilizing the sepsis score and blood culture status to distinguish foals into categories (healthy, sick non-septic and sepsis) has inherent limitations and low reported sensitivities in neonatal foals [3,7,8,20,36]. A foal may have a low SS but be blood culture positive with a non-pathogenic isolate or contaminant but be classified as septic. False positive blood cultures have been documented in the human literature and have been associated with increased hospital stays, inappropriate administration of antimicrobials and unnecessary hospital expense [37–39]. Blood cultures yielding non-pathogenic isolates can also influence sepsis scoring systems and categorization schemes [38,40,41]. Similarly, a foal might have a SS of 6 to 11 and be culture negative but later goes on to develop clinical and clinicopathologic derangements consistent with sepsis. This subject would be falsely classified as sick non-septic using the current classification criteria. Several reasons explaining negative blood cultures in septic foals may include the fastidious nature of some organisms that do not grow well in culture media or low circulating numbers of bacteria in the bloodstream at the time of sampling [3]. Prior administration of antimicrobials at the time of sample collection could also result in a negative culture result. This can certainly be true of foals with more severe clinical signs and a higher suspicion for neonatal sepsis. It is common protocol for ambulatory veterinarians to administer foals antimicrobials and plasma prior to referral when there is a high index of suspicion for infection. Antimicrobials may kill or inhibit growth of bacteria and plasma can opsonize bacteria in the sample [34,42]. There is strong evidence in humans to begin antimicrobials immediately on the suspicion of sepsis.

Guidelines in human medicine report that a delay in treatment is associated with increased morbidity and mortality [1,43,44]. The classification scheme used in this study has a sensitivity of 60% and specificity of 61% with an area under the curve of 0.71 [19]. The additional classification criteria of a positive blood culture possesses a documented sensitivity of only 61.9% in septic human neonates [6]. Given these findings, further study is needed to more accurately classify foals, particularly septic foals versus bacteremic foals.

Changing trends in cultured organisms have emphasized the need for a reliable method of sepsis detection and bacteremia in the context of therapeutic decision making. In choosing antimicrobials, spectrum is heavily considered. Our data illustrate an overall increase prevalence of Gram-positive organisms. 100% of foals that did not survive to discharge in this study cultured at least one Gram-positive organism. The general increase in Gram-positive isolates is consistent with recent reports identifying a shift away from predominantly Gram-negative isolates [45,46]. Gram-positive isolates have shown a lack of predictability in their antimicrobial susceptibility patterns [45,46]. Extensive administration of antimicrobials has placed selection pressures on isolates presumably favoring the growth of resistant species and contributing to the need for effective bacterial identification and sepsis diagnostics.

In conclusion, plasma cell-free DNA concentration levels appear to be inadequate as a diagnostic marker for foals with sepsis or biomarker to predict survival in a neonatal population. Similar to studies in adult humans and dogs, the utility of cfDNA warrants further study in adult horses, notably horses with critical illness where cfDNA may have prognostic value in an immunologically mature subject but is yet to be determined. Further, pursuit of a robust and accurate method to diagnose sepsis is needed as illness severity scoring systems (e.g. sepsis score) and one-time blood culture methods appear woefully imprecise in this population.

## Acknowledgements

The authors wish to thank the interns, residents and nurses at New Bolton Center for sample collection and their dedication to the management of the foals in this study.

## REFERENCES

1. Fielding CL, Magdesian KG. Sepsis and Septic Shock in the Equine Neonate. Vet Clin North Am. 2015;31:483–496.

2. Paradis, MR. Vet Clin North Am Equine Pract. Equine Practice. 1994;10:109–135.

3. Giguere S, Weber EJ, Sanchez LC. Factors associated with outcome and gradual improvement in survival over time in 1065 equine neonates admitted to an intensive care unit. Equine Vet J. 2017;49:45–50.

4. Hurcombe SD, Toribio RE, Slovis N. Blood arginine vasopressin, adrenocorticotropin hormone and cortisol concentrations at admission in septic and critically ill foals and their association with survival. J Vet Intern Med. 2008;22:639–647.

5. Hurcombe SD, Toribio RE. Slovis NM. Calcium regulating hormones and serum calcium and magnesium concentrations in septic and critically ill foals and their association with survival. J Vet Intern Med. 2009;23:335–343.

6. Weinbren MJ, Collins M, Heathcote R, Umar M, Nisar M, Ainger C, et al. Optimization of the Blood Culture Pathway: A Template for Improved Sepsis Management and Diagnostic Antimicrobial Stewardship. J Hosp Infect. 2018;98:232–235.

7. Pusterla N, Mapes S, Burne BA, Magdesian KG. Detection of blood stream infection in neonatal foals with suspected sepsis using real-time PCR. Vet Rec. 2009;165:114–117.

8. Hackett ES, Lunn DP, Ferris RA, Horohov DW, Lappin MR, McCue PM. Detection of bacteremia and host response in healthy neonatal foals. Equine Vet J. 2015;47:405–409.

9. Corley KT, Furr MO. Evaluations of a score designed to predict sepsis in foals. J Vet Emerg Crit Care. 2003;13:149–155.

10. Letendre JA, Goggs R. Determining prognosis in canine sepsis by bedside measurement of cell-free DNA and nucleosomes. J Vet Emerg Crit Care. 2018;28:503–511.

11. Avriel A, Wiessman MP, Almog Y, Perl Y, Novack V, Galante O et al. Admission Cell Free DNA Levels Predict 28-day Mortality in Patients with Severe Sepsis in Intensive Care. PLoS ONE. 2014;9:e100514 1–7.

12. Rhodes A, Wort SJ, Thomas H, Wort SJ, Thomas H, Collinson P et al. Plasma DNA concentration as a predictor of mortality and sepsis in critically ill patients. Crit Care. 2006;10:R60.

13. Yipp BG, Kubes P. NETosis: how vital is it? Blood. 2013;122:2784–2794.

14. Streleckiene G, Forster M, Inciuraite R, Lukosevicius R, Skieceviciene J. Effects of Quantification Methods, Isolation Kits, Plasma Biobanking, and Hemolysis on Cell-Free DNA Analysis in Plasma. Biopreserv Biobank. 2019;17: doi: 10.1089/bio.2019.0026.

15. Dragan AI, Casas-Finet JR, Bishop ES, Strouse RJ, Schenerman MA, Geddes CD. Characterization of PicoGreen Interaction with dsDNA and the Origin of Its Fluorescence Enhancement Upon Binding. Biophys J. 2010;99:3010–3019.

16. Li X, Yuhua Wu, Zhang Li, Cao Y, Li Y, Li J et al. Comparison of three common DNA concentration measurement methods. Anal Biochem. 2014;451:18–24.

17. Rhodes A, Cecconi M. Cell-free DNA and outcomes in sepsis. Crit Care 2012;16:170.

18. Saukkonen K, Lakkisto P, Pettila V, Varpula M, Karlsson S, Ruokonen E et al. Cell-free plasma DNA as a predictor of outcome in severe sepsis and septic shock. Clin Chem. 2008;45:1000–1007.

19. Letendre KA, Goggs R. Measurement of plasma cell-free DNA concentrations in dogs with sepsis, trauma and neoplasia. J Vet Emerg Crit Care. 2017;3:307–314.

20. Wong DM, Ruby RE, Dembek KA, Barr BS, Ruess SM, Magdesian KG et al. Evaluation of updated sepsis scoring systems and systemic inflammatory response syndrome criteria and their association with sepsis in equine neonates. J Vet Intern Med. 2018;32:1185–1193.

21. Dembek KA, Onasch K, Hurcombe SDA, MacGillivray KC, Slovis NM, Barr BS et al. Renin-Angiotensin-Aldosterone system and hypothalamic-pituitary-adrenal axis in hospitalized neonatal foals. J Vet Intern Med. 2013;27:331–338.

22. Barnsick RJIM, Hurcombe SDA, Dembek KA, Frazer ML, Slovis NM, Saville WJA, et al. Somatotropic axis resistance and ghrelin in critically ill foals. Equine Vet J. 2014;46:45–49.

23. Sato A, Nakashhima C, Abe T, Kato J, Hirai M, Nakamura T et al. Investigation of appropriate pre-analytical procedure for circulating free DNA from liquid biopsy. Oncotarget. 2018;9:31904–31914.

24. Hollis A, Wilkins P, Palmer J, Boston RC. Bacteremia in equine neonatal diarrhea: a retrospective study (1990-2007). J Vet Intern Med. 2008;22:1203–1209

25. Tagawa M, Shimbo G, Inokuma H, Miyahara K. Quantification of plasma cell-free DNA levels in dogs with various tumors. J Vet Diagn Invest. 2019;31(6):836–843.

26. Jeffery U, Kimura K, Gray R, Lueth P, Bellaire B, LeVine D. Dogs cast NETs too: Canine neutrophil extracellular traps in health and immune-mediated hemolytic anemia. Vet Immunol Immunopathol. 2015;168:262–268.

27. Rushton JG, Ertl R, Klein D, Tichy A, Nell B. Circulating cell-free DNA does not harbour a diagnostic benefit in cats with feline diffuse iris melanomas. J Feline Med Surg. 2019;21(2):124–132.

28. Huang T, Yang Z, Chen S, Chen J. Predictive value of plasma cell-free DNA for prognosis of sepsis. Zhonghua Wei Zhong Bing Ji Jiu Yi Xue. 2018;30(10):925–928.

29. Tuaeva NO, Abramova ZI, Softonov VV. The Origin of Elevated Levels of Circulating DNA in Blood Plasma of Premature Neonates. Ann N Y Acad Sci. 2008;1137:27–30.

30. Perkins GA, Wagner B. The development of equine immunity: Current knowledge on immunity in the young horse. Eq Vet J. 2014;47:267–274.

31. Aucamp J, Bronkhorst AJ, Badenhorst CPS, Pretorius PJ. The diverse origins of circulating cell-free DNA in the human body: a critical re-evaluation of the literature. Biol Rev Camb Philos Soc. 2018;93:1649–1683.

32. Goggs R, Jeffery U, LeVine DN, Li RHL. Neutrophil-Extracellular Traps, Cell-Free DNA, and Immunothrombosis in Companion Animals: A Review. Vet Pathol. 2020;57(1):6–23.

33. Li, HL, Ng, G, Tablin, F. Lipopolysaccharide-induced neutrophil extracellular trap formation in canine neutrophils is dependent on histone H3 citrullination by peptidylarginine deiminase. Vet Immunol Immunopathol. 2017;193–194:29–37.

34. McTaggart A, Yovich JV, Penhale J, Raidal SL. A comparison of foal and adult horse neutrophil function using flow cytometric techniques. Res Vet Sci. 2001;71(1):73–79.

35. Borchers A, Wilkins PA, Marsh PM, Axon JE, Read J, Castagnetti C. et al. Association on admission L-lactate concentration in hospitalized equine neonates and with presenting complaint, periparturient events, clinical diagnosis and outcome: a prospective multicenter study. Equine Vet J. 2012;44(41):57–63.

36. Furr M, McKenzie H. Factors associated with the risk of positive blood culture in neonatal foals presented to a referral center (2000-2014). J Vet Intern Med. 2020;1–13.

37. Norberg A, Christopher NC, Ramundo ML, Bower JR Berman SA. Contamination rates of blood cultures obtained by dedicated phlebotomy vs intravenous catheter. JAMA. 2003;289:726–729.

38. Weinbaum FL, Lavie S, Danek M, Sixsmith D, Heinrich GF, Mills SS. Doing it right the first time: quality improvement and the contaminant blood culture. J Clin Microbiol. 1997;35:563–565.

39. Hall KK, Lyman JA. 2006. Updated Review of Blood Culture Contamination. Clin Microbiol Rev. 2006;19:788–802.

40. Bates DW, Goldman L, Lee TH. Contaminant blood cultures and resource utilization: the true consequences of false-positive results. JAMA. 1991;265:365–369.

41. Dunagan WC, Woodward RS, Medoff G, Gray JL III, Casabr E, Smith MD et al. Antimicrobial misuse in patients with positive blood cultures. Am J Med. 1989;87:253–259.

42. Scheer CS, Fuchs C, Grundling M, Vollmer M, Bast J, Bohnert JA et al. Impact of antibiotic administration on blood culture positivity at the beginning of sepsis: a prospective clinical cohort study. Clin Micribiol Infect. 2019;25:326–331.

43. Weiss SL, Fitzgerald JC, Balamuth F, Alpern ER, Lavelle J, Chilutti M et al. Delayed antimicrobial therapy increases mortality and organ dysfunction in pediatric sepsis. Crit Care Med. 2014;42:2409–2417.

44. Wisdom A, Eaton V, Gordon D, Daniel S, Woodman R, Phillips C. INITAIT-E.D.: impact of timing of INITIation of Antibiotic Therapy on mortality of patients presenting to an Emergency Department with sepsis. Emerg Med Australas. 2015;27:196–201.

45. Theelan MJP, Wilson WD, Edman JM, Magdesian KG, Kass PH. Temporal trends in prevalence of bacteria isolated from foals with sepsis: 1979-2010. Equine Vet J. 2014;46(2):169–73.

46. Russell CM, Axon JE, Bilshen A, Begg AP. Blood culture isolates and antimicrobial sensitivities from 427 critically ill neonatal foals. Aust Vet J. 2008:86(7):266–271.

